# A Robust Proteomics-Based Method for Identifying Preferred Protein Targets of Synthetic Glycosaminoglycan Mimetics

**DOI:** 10.1101/2025.01.23.634492

**Authors:** Daniel K. Afosah, Ravikumar Ongolu, Rawan M. Fayyad, Adam Hawkridge, Umesh R. Desai

## Abstract

A robust technology is critically needed for identifying preferred protein targets of glycosaminoglycans (GAGs), and synthetic mimetics thereof, in biological milieu. We present a robust 10-step strategy for identification and validation of preferred protein targets of highly sulfated, synthetic, small, GAG-like molecules using diazirine-based photoaffinity labeling– proteomics approach. Our work reveals that optimally designed, homogeneous probes based on minimalistic photoactivation and affinity pulldown groups coupled with rigorous proteomics, biochemical and orthogonal validation steps offer excellent potential to identify preferred targets of GAG mimetics from the potentially numerous possible targets that cloud GAG interaction studies. Application of this 10-step strategy for a promising highly sulfated, small GAG mimetic led to identification of only a handful of preferred targets in human plasma. This new robust strategy will greatly aid drug discovery and development efforts involving GAG sequences, or sulfated small mimetics thereof, as leads.

## INTRODUCTION

Sulfated glycosaminoglycans (GAGs), nature’s eclectic biopolymers, are unique in terms of both their structure and function. Structurally, GAGs are the most diverse biopolymers with the number of unique sequences exceeding 10^9^ for polymeric chains as small as 6 units long.^1-3^ Functionally, GAGs impact a large number of fundamental processes such as out-in signal transduction, cell metabolism, cell growth/differentiation, cell–cell and cell–matrix communication, etc.^4-7^ These diverse impacts arise from their interaction with more than 3,000 proteins, which represents the size of the current GAG interactome.^4,6^ Despite this massive interactome, except for heparins that target antithrombin, no GAG-like molecule has reached the clinic as yet. This implies that the huge drug discovery potential offered by the known GAG interactome remains largely untapped.

A key challenge to discovering GAG-like drugs is identifying their preferred targets in complex biological settings. Several technologies have been explored for this purpose, including *in silico* library screening,^8^ microarray screening,^7,9,10^ and biophysics-based high-throughput screening.^11,12^ We reasoned tandem photoaffinity labeling (PAL) and proteomics experimentation promises a more powerful approach to identify preferred protein target(s) of GAGs. Yet, the application of PAL–proteomics technology remains undeveloped for highly sulfated GAG-like small molecules. A key challenge in realizing the promise of PAL–proteomics technology is overcoming the numerous non-specific, electrostatic interactions that typically cloud and distort the results.

Recently, Hsieh-Wilson and colleagues applied PAL-proteomics to chondroitin sulfate E (CS-E) polymers, wherein 54 proteins were identified as likely targets.^13^ The authors used non-selective placement of a benzophenone-based photoactivatable group along the CS-E polymeric chain, which although successful could impact target potency and fidelity. Considering that GAG-binding pockets on proteins are shallow coupled with largely moderate-affinity GAG-protein interactions,^8,14,15^ a structurally homogeneous PAL probe is likely to be critical. In fact, early applications of PAL technology to GAGs, such as heparins, have suffered from probe heterogeneity, target potency and probably target selectivity.^1,8,14^ In turn, probe heterogeneity introduces inefficient and heterogeneous photolabeling,^16^ challenging target pulldown and characterization,^5^ and other issues.^7^

A robust PAL-proteomics technology is critically needed for GAGs, and mimetics thereof, for advancing drug discovery and development efforts. *A priori*, this technology would not only pinpoint preferred targets of highly sulfated small molecules in biological milieu but may also offer early alerts on off-target effects. In this work, we present a robust strategy for identification and validation of the preferred targets of highly sulfated small GAG-like molecules using diazirine-based PAL-proteomics technology. Our work shows that designing a homogeneous probe that minimally alters the parent structure coupled with rigorous application of proteomics can identify preferred protein targets of GAG mimetics, i.e., highly sulfated small molecules. We present a 10-step strategy that can be implemented using a promising anti-cancer synthetic GAG mimetic G2.2 (**Figure 1A**) as an example, which rapidly identified only a few high-affinity targets from the hundreds present in human plasma and dozens labeled by the photoaffinity probe. Our strategy is likely to be particularly helpful in early drug discovery efforts of GAGs and GAG-like small molecules.

**FIGURE 1.**
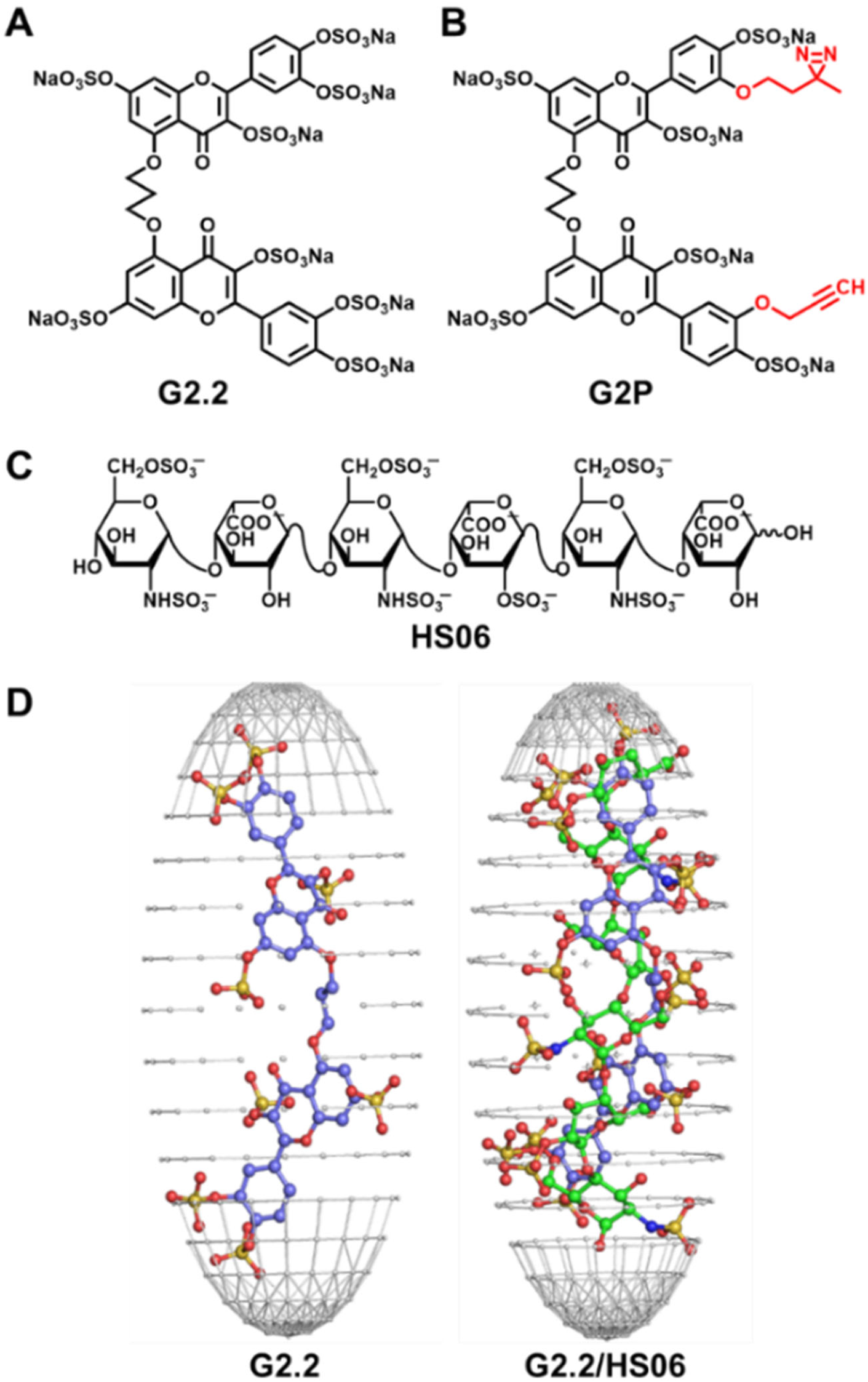
Structures of synthetic GAG mimetic G2.2 (A) and its photoaffinity probe G2P (B) containing diazirine and alkyne moieties (shown in red). (C) Molecular models of G2.2 (left) and G2.2 (blue) overlaid on HS06 (green) with sulfates shown in atom-type color. (D) Structure of HS06, which is functionally and structurally mimicked by G2.2.

## RESULTS AND DISCUSSION

### Rationale and Design of PAL Probe

Over the past decade, we have developed the field of <u>n</u>on-<u>s</u>accharide <u>G</u>AG <u>m</u>imetics (NSGMs), which are homogeneous, synthetic, highly sulfated, aromatic-scaffold based small molecules that mimic the function of natural GAGs.^17,18^ NSGMs typically bind in the electropositive and shallow GAG-binding pocket of proteins and conformationally alter their biological function.^19-21^ In fact, highly promising drug-like GAG mimetics have been designed as anti-thrombotics,^19,20^ anti-cancer,^22,23^ anti-inflammatory^24^ and anti-infectives^25^ with minimal toxicity. To advance these molecules to the clinic, target selectivity under physiologic/pathologic conditions is important to establish, especially because GAGs are known to engage thousands of proteins.^4,6^ Hence, we sought to develop PAL-proteomics technology in identifying the preferred target(s) (and possible off-target effects) of these NSGMs in an unbiased manner.

Within our library of more than 100 NSGMs is a promising mimetic G2.2 (**Figure 1A**). G2.2 effectively mimics heparan sulfate hexasaccharide (HS06) exhibits highly selective anti-cancer stem cell activity.^17-19,23^ The preferred target(s) of G2.2 on cancer stem cells remains to be identified. Although PAL-proteomics technology should in principle be very useful for such a goal, it remains poorly developed for GAGs. Hence, we designed photoaffinity probe G2P (**Figure 1B**), which placed emphasis on introducing minimal changes to the parent molecule. The critical diazirine and alkyne groups were placed at the two 3’-positions, which have been shown earlier to retain anti-cancer activity.^19,23^ Furthermore, we chose the diazirine group (<2Å), instead of the benzophenone group (∼10Å) suggested by the Hsieh-Wilson group,^13^ for its much small footprint. We reasoned that target selectivity of a small sulfated GAG-like molecule could be significantly impacted by the bulky benzophenone. We also used small alkyl linkers to attach diazirine and alkyne groups to keep G2P as close to G2.2 as possible.

### Synthesis of G2P

The photoaffinity probe of G2.2 was synthesized in eight steps from quercetin by utilizing the hydrogen-bonding capabilities of its 5- and 3’ hydroxy groups (**Figure 2A**, see also **Scheme S1** for details). Briefly, partial MOM-protection^23,26^ of quercetin gave key intermediate **2**, which was alkynylated or diazirinylated to yield the two halves of G2P, i.e., **3** and **4** (see Supplementary Information for details). These were then coupled using a difunctional linker to give precursor **5**, which was deprotected and sulfated under microwave conditions^26,27^ followed by Na^+^-exchange chromatography to yield G2P in high purity (see **Figure S1**).

**FIGURE 2.**
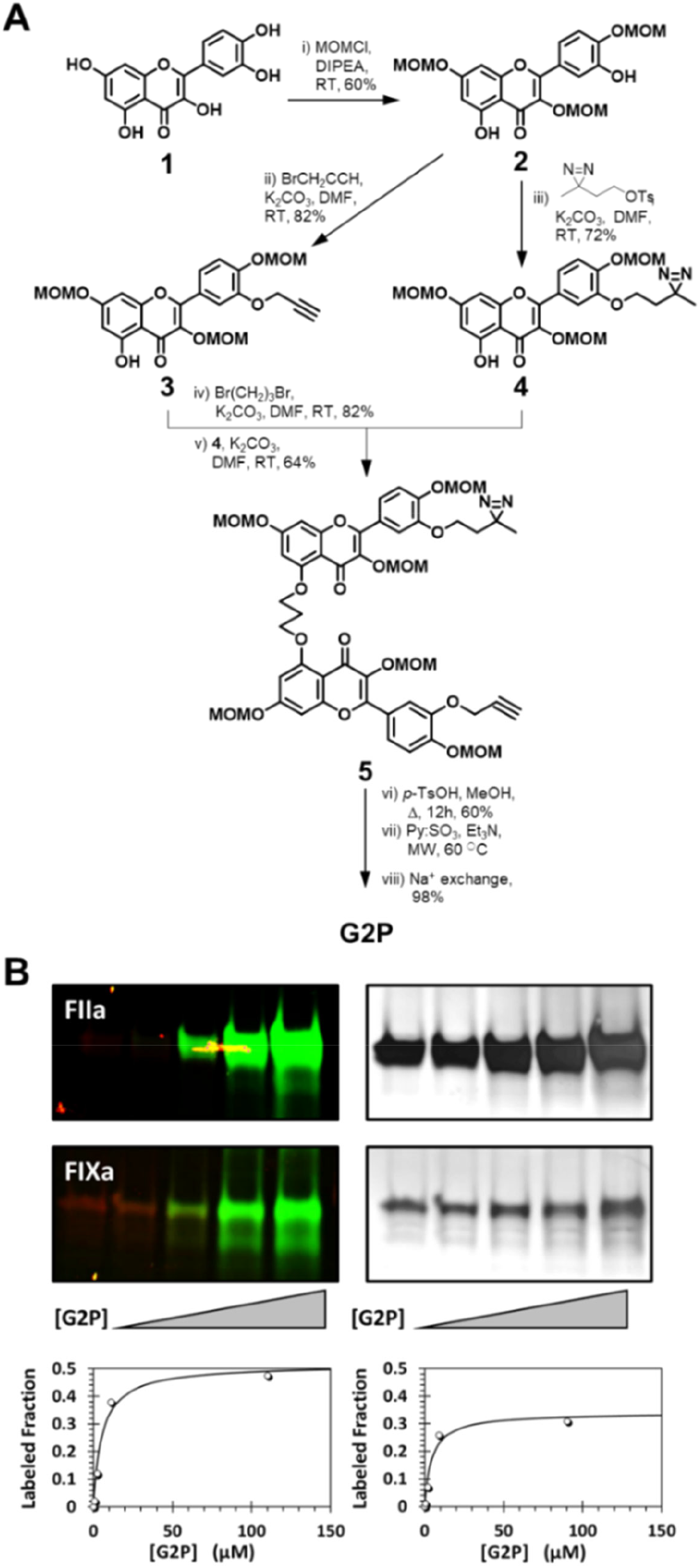
(A) Synthesis of G2P from quercetin (1) in eight steps. (B) Binding and photolabeling property of G2P against two test proteases, thrombin (FIIa) and factor IXa (FIXa), of the coagulation cascade. Increasing levels of G2P levels resulted in higher photoaffinity labeling (fluorescence gels) as a proportion of the total protein (silver stained gels), which can be analyzed as hyperbolas for affinity calculations (graphs).

### G2P photolabels known GAG-binding proteins in a classic affinity-dependent manner

To assess the applicability of G2P, we first studied its binding to thrombin, which is well-established prototypic, non-selective target of all highly sulfated GAGs and GAG mimetics.^18,28^ A fixed concentration (2 µM) was incubated with G2P (0→100 µM) for 10 min, photolysed (λ=368 nm) for 30 min at 4°C, “click”-coupled with rhodamine azide, and electrophoresed (**Table S1**). Fluorescence imaging revealed excellent efficiency (∼50%) of photolabeling (**Figure 2B**), which was estimated from the intensity of silver stained gels. In fact, a characteristic hyperbolic profile of the fraction labeled was observed, which implied that G2P binds and covalently modifies thrombin in a classic affinity-dependent manner. This raises the hypothesis that PAL experimentation could be performed to preferentially label proteins in a mixture.

We also tested G2P against another coagulation factor, FIXa, which is also a known target of sulfated polysaccharides,^29^ and observed equivalent results (**Figure 2B**). These results are significantly different from those with diazirine-labeled CS-E, which was found to be inefficient.^13^ Although the exact reason for this difference is not obvious, we predict that the proximity of the diazirine group to the GAG scaffold and the homogeneity of the PAL probe are critical.

We next assessed the applicability of G2P in identifying protein targets in a biological mixture. A mixture of 10 proteins (hemoglobin, heparin cofactor II, prothrombin, factor IX, antithrombin, factor X, factor XI, IGF1, factor XIII, and bovine serum albumin) was incubated with G2P, irradiated and covalently clicked with rhodamine (**Table S2**). As observed for FIIa and FIXa, excellent photolabeling was observed (**Figure S2**). Several control reactions, e.g., exclusion of the probe, UV irradiation or the fluorophore tag, ensured high rigor in photolabeling. Also, photo-cross-linking was severely impaired in the presence of parent G2.2 as a competitor. Interestingly, preferential photolabeling was observed revealing early support for the affinity-based hypothesis presented above.

### Strategy for Identifying Preferred Protein Targets in a Biological Milieu

We first evaluated photolabeling of pooled human plasma in its native form and numerous challenges arising from the presence of high abundance proteins, which overwhelm the proteomics results (not shown). Using protein depletion columns (Pierce™ Top-12) that reduce the levels of the most abundant, followed by SDS-PAGE analysis revealed that only a subset of proteins was photolabeled (**Figure S3**). Absence of G2P or UV irradiation, or competition with parent G2.2, led to concomitant decreases in photolabeling supporting applicability of the technology to biological setting.

Next, we developed a 10-step algorithm to identify preferred target(s) of G2P (**Figure 3**). First, pooled human plasma depleted of its high abundance proteins was treated with several concentrations of G2P (0→75 µM), UV irradiated, reacted with biotin-PEG-azide (or rhodamine-azide for gel imaging, if needed). The biotinylated product was resolved into streptavidin-bound and unbound fractions, which upon PAGE analysis revealed bands in the latter that increased in tandem with G2P levels (**Figure S4**). The bound fraction (Bf) at each G2P concentration (0, 7.5, 15, 30 and 75 µM) was studied in multiple proteomics runs (n≥3), in which an identical amount of each sample was trypsinized and subjected to LC-MS/MS analysis.

**FIGURE 3.**
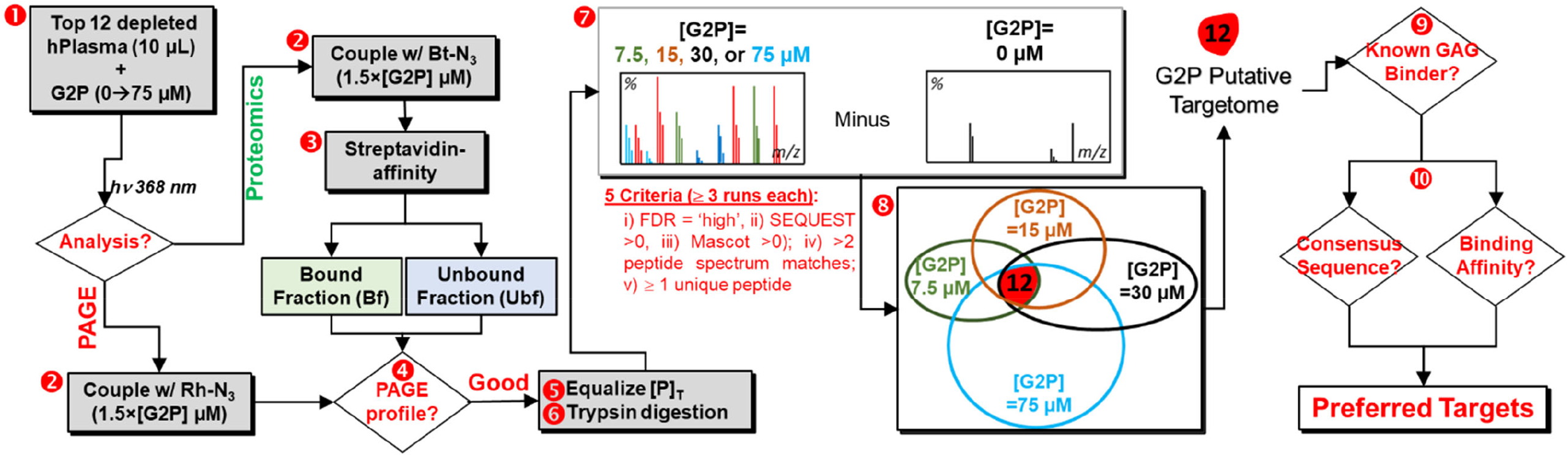
The 10-step strategy based on PAL-proteomics to identify the preferred targets of G2P, and thereby G2.2, in human plasma. Steps 1 – 4 involve PAL methods, steps 5 – 8 correspond to proteomics, and steps 9 & 10 involve validation of targets. This strategy rests on rigorous application of steps 7, 8 and 10, which focus on identifying the ‘preferred’ targets. The human plasma G2P targetome was reliably identified to contain 12 proteins (step 8) from >80 identified as plausible targets. See text for details.

We used several stringent criteria within Proteome Discover, Mascot, SEQUEST and Byonic packages used for proteomics analysis to ensure high-fidelity output. These included i) a combined false discovery rate (FDR) set to high confidence; ii) Mascot score >0; iii) SEQUEST score >0; iv) at least two peptide spectrum matches; and v) identification of at least one unique peptide for each protein identified as a putative target (see Supplementary Information). Together, these criteria help wade out false positives, which usually abound and occlude identification of true positives, especially for proteins binding to GAG-like molecules. Next, we implemented a biochemical filter. Proteins identified in the control ([G2P]=0 µM) cannot be classified preferred targets and hence were excluded (Step 7, **Figure 3**). This filter resulted in putative protein targets ranging from 22–80 for G2P concentrations in the range of 7.5–75 µM. Finally, only those proteins that were common to all experimental conditions (i.e., 7.5–75 µM G2P) were deemed ‘preferred’ G2P targets. This led to 12 proteins as the preferred G2P, and potentially G2.2, protein ‘targetome’ (step 8, **Figure 3**).

To confirm the preferred targets, we compared the 12 protein G2P targetome (see **Table S5** for the list) with the published database of GAG-binding proteins.^4^ Ten of the 12 (∼75%) had been known to bind GAGs (Step 9, **Figure 3**), whereas two were new. An orthogonal prediction of GAG binding is the presence of Cardin-Weintraub (CW) motifs (‘XBBXBX’ and ‘XBBBXXBX’ sequences).^8,30^ One or more CW motifs were present in ten proteins (**Figure 4A**). These two predictive methods implied that our PAL-proteomics results were largely supported.

**FIGURE 4.**
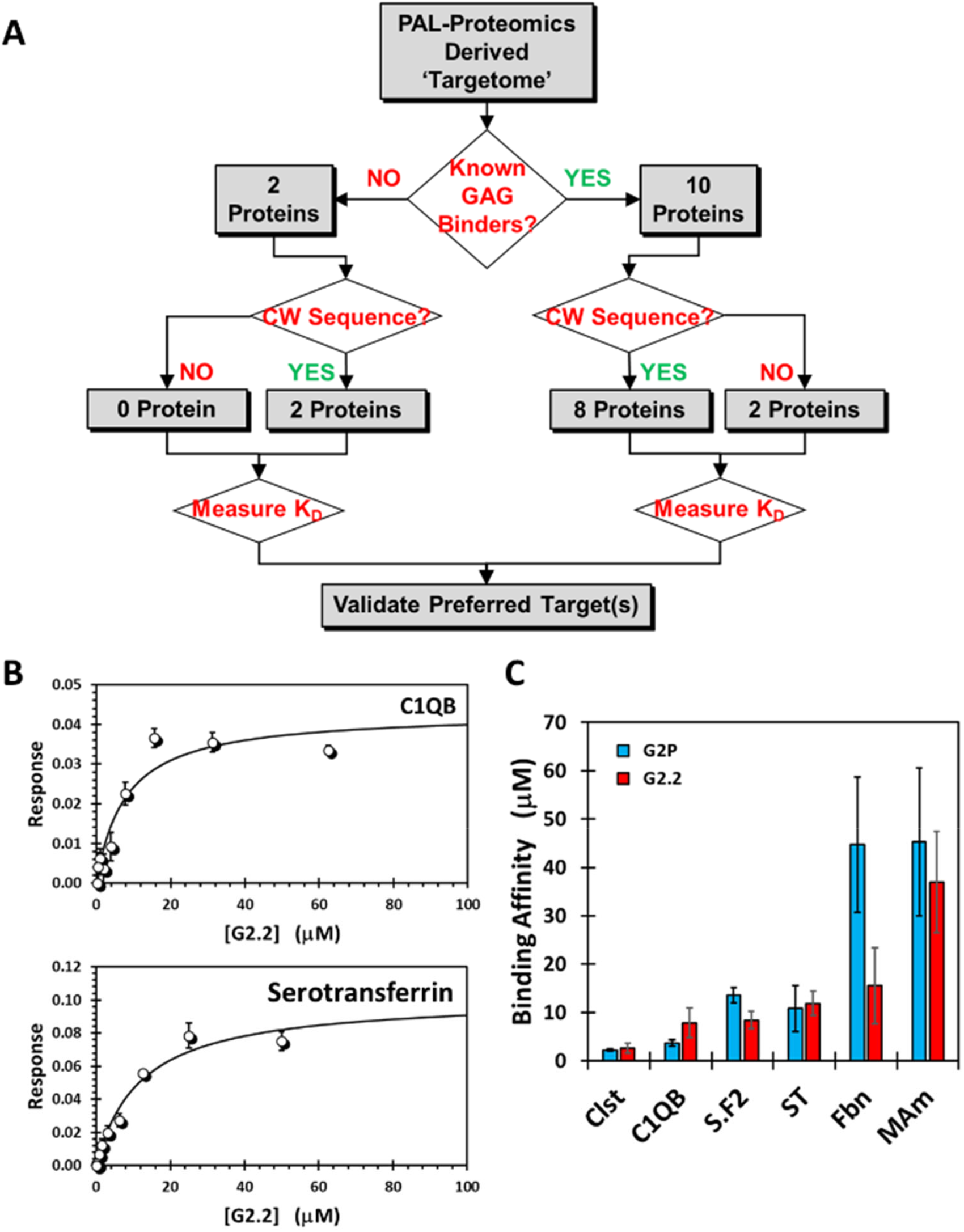
(A) Distribution of proteins identified as preferred targets of G2P using literature reports and Cardin-Weintraub (CW) consensus sequence analysis. See also Table S5 for details. (B) Binding affinities of G2.2 for two of the representative six protein targets identified as preferred ones through PAL-proteomics. His-tagged proteins, labeled with RED-tris-NTA dye, were used in microscale thermophoresis (MST)-based titrations to measure *K*_D_ of interaction. (C) Comparison MST-measured *K*_D_s for G2.2 (red) and G2P (cyan). Error bars show ±1 SE. See also Figures S5 and S6 for all profiles.

Yet, predictions based on polymeric GAGs, which rely on their chain length, may not stand for small molecules. Hence, we measured the thermodynamic affinities of representative proteins for G2.2. We selected six proteins considering the GAG database/CW motifs predictions as well as possible role of G2.2 in cancer biology. Of these, clusterin, complement C1Q (C1QB), fibronectin, and serpin F2 were known to bind GAGs, whereas serotransferrin and *N*-acetyl muramoyl-*L*-alanine amidase (NAm) were not known to bind to GAGs. In each case, characteristic microscale thermophoresis (MST) profiles were observed for G2.2 binding to recombinant proteins, which when analyzed using stoichiometric binding isotherms yielded affinities of 2.6, 7.8, 8.4, 11.9, 15.5, and 36.9 µM for clusterin, C1QB, serotransferrin, serpin F2, fibronectin, and NAm, respectively (**Figure 4B**, see also **Figure S5** for all profiles). These results confirm the identification of the preferred targets of G2.2. Interestingly, two of the three most potent targets, i.e., serotransferrin and C1QB, play important roles in cancer biology.^31-35^ The identification of serotransferrin as one of the preferred targets of anti-cancer stem cell agent G2.2 is even more relevant considering serotransferrin’s role in cancer stem cell biology.^31-33^

A key question arises whether G2P is a robust PAL probe for G2.2. To answer this question, we measured G2P affinities for the six proteins using the same technique. **Figure 4C** compiles the results. In effect, the affinities of G2P and G2.2 are highly comparable and change in tandem, except perhaps for fibronectin (44.7±13.9 and 15.5±7.8 µM, respectively; see **Figure S6**). Overall the results validate the 10-step algorithm for elucidating preferred protein targets of highly sulfated, small GAG mimetics through a diazirine-based PAL-proteomics technology.

## CONCLUSIONS

The literature presents thousands of proteins as possible targets of GAGs, and therefore of their mimetics too.^1-7^ Majority of these proteins are not likely to be relevant targets. Unfortunately, few technologies are available that can rapidly and conclusively identify preferred targets. Although PAL–proteomics technology is theoretically very promising,^36,37^ especially for small hydrophobic drug-like molecules (e.g., such as doxorubicin,^38^ danazol^39^ and cis-platin,^40^), it has been very difficult to implement it for GAG-like molecules. This report presents the first PAL– proteomics algorithm for synthetic, drug-like, small GAG mimetics.

The success of our PAL–proteomics strategy originates from some key structural and algorithmic design principles. First, our PAL probe was homogeneous. Second, probe elements (diazirine and alkyne) were designed to ensure least disruption of the parent GAG agent. Third, the probe design emphasized minimalistic features (alkyl linkers). Fourth, our 10-step proteomics algorithm emphasized reproducibility and biochemical principles. Finally, orthogonal validation steps enhanced rigor. We believe that these fundamental points are can be implemented for nearly all defined GAG sequences and mimetics thereof. Thus, this PAL–proteomics technology is broad applicable.

While broadly implementable, our technology does not mute all the challenges. At the structural level, success relies on probe design, which is dependent on knowledge about structure–activity relationship of the parent GAG/mimetic. At the biochemistry level, the technology is expected to works best when non-specific competition, arising from abundant proteins, is minimized. At the proteomics level, it is important to ascertain that the risk of false negatives is low.

A byproduct of this work is identification of putative targets of G2.2 in human plasma. G2.2 is a highly promising inhibitor of cancer stem cells^22,23,31^ and mimics HS06 (**Figure 1D**),^17^ which has recently been found to inhibit cell-surface bound insulin-like growth factor–1 receptor (IGF-1R).^41^ Thus, G2.2 may target IGF-1R on cancer cells, while also engaging the 12 preferred plasma proteins. It remains to be clarified whether G2.2 would prefer cancer cell surface IGF-1R or its human plasma targets. This PAL–proteomics technology is expected to be particularly useful for such an unbiased endeavor.

Overall, our PAL–proteomics strategy is expected to be particularly useful in drug discovery and development efforts involving GAG sequences, or sulfated small mimetics thereof, as leads. The technology is likely to find particular use in identifying on and off targets.

## Abbreviations

Bf: bound fraction
CW: Cardin-Weintraub motifs
GAGs: glycosaminoglycans
Hp: heparin
HS: heparan sulfate
HS06: heparan sulfate hexasaccharide
NSGMs: non-saccharide glycosaminoglycan mimetics
PAL: photoaffinity labeling

## Supporting Information

Methods describing synthesis and characterization of G2P; Proteomics experimentation and analysis; Scheme S1, Figures S1 – S6 and Tables S1 – S5. Also available are raw proteomics datafiles, which can be downloaded from the publisher’s website.

## Author Contributions

DKA and RKO performed experiments, analyzed data, and wrote the first draft of manuscript. RF performed RP-IP UPLC-MS experimentation; AH helped with initial proteomics experimentation design. URD analyzed data, prepared figures, and revised the manuscript. All authors have given approval to the final version of the manuscript.

## Acknowledgements

We thank W. Keith Ray of the Virginia Polytechnic Mass Spectrometry Incubator for performing the proteomics work.

## TABLE OF CONTENT GRAPHIC

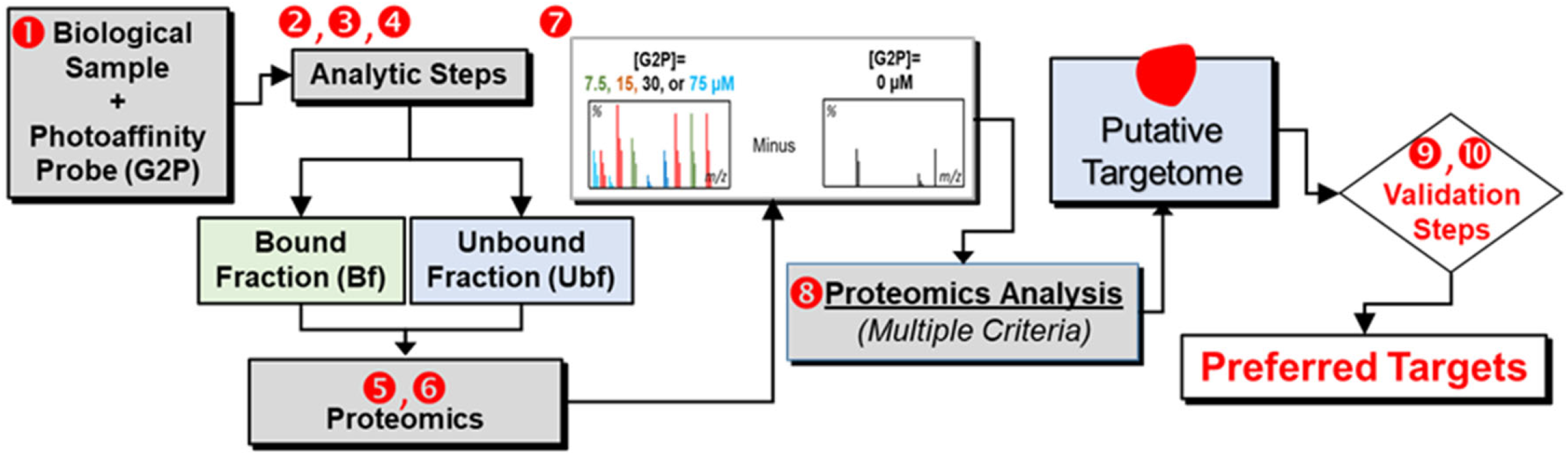

