## Supplementary material for "A Robust Proteomics-Based Method for Identifying Preferred Protein Targets of Synthetic Glycosaminoglycan Mimetics": SupplInfo



| # | Content | Pg |
|---|---|---|
| 1 | Chemicals, reagents, and proteins | S2 |
| 2 | Scheme S1. Synthesis of G2P | S3 |
| 3 | Experimental methods: Synthesis and characterization of G2P | S4 |
| 4 | Figure S1. UPLC-MS profile of G2P | S6 |
| 5 | Experimental methods: Photoaffinity labeling of proteins in buffer | S7 |
| 6 | Table S1. Concentrations used in G2P-labeling of FIIa and FIXa | S7 |
| 7 | Table S2. Conditions for G2P-labeling of protein mix | S7 |
| 8 | Figure S2. PAGE analysis of photolabeled mixture of proteins | S8 |
| 9 | Experimental methods: G2P photolabeling of human plasma proteins | S9 |
| 10 | Table S3. Conditions for G2P-labeling of human plasma proteins | S9 |
| 11 | Figure S3. PAGE analysis of photolabeled human plasma proteins | S10 |
| 12 | Experimental methods: Affinity pull-down G2P-labeled human plasma proteins | S11 |
| 13 | Table S4. Conditions for G2P-labeling and affinity pulldown | S11 |
| 14 | Figure S4. PAGE of fractions following G2P-labeling and affinity pulldown | S12 |
| 15 | Experimental methods: Proteomics – preparation, experimentation, and analysis | S13 |
| 16 | Experimental methods: MST of G2.2 and G2P binding to protein targets | S14 |
| 17 | Table S5. Prediction on GAG-binding property of 12 preferred targets | S15 |
| 18 | Figure S5. Affinity of G2P for representative targets | S16 |
| 19 | Figure S6. Affinity of G2.2 for representative targets | S17 |

**Chemicals, reagents, and proteins**—All chemicals and anhydrous organic solvents were purchased from either Sigma-Aldrich (Milwaukee, WI) or Fisher (Pittsburgh, PA), unless otherwise stated. Other solvents used were of reagent gradient and used as purchased. Analytical TLC was performed using UNIPLATE™ silica gel GHLF 250 um pre-coated plates (ANALTECH, Newark, DE). Silica gel (200-400 mesh, 60 Å), azide-fluor 488 (AF488), TBTA, TCEP, $CuSO_4$, and unfractionated heparin (UFH) were from Sigma-Aldrich. Biotin, azido-PEG3-biotin conjugate, bovine serum albumin (BSA), and Pierce streptavidin agarose resin were from Fisher. Pierce silver staining kit and depletion spin columns for abundant proteins were from ThermoFisher Scientific. SDS-PAGE gels, Tris/glycine/SDS running buffer and Laemmli sample buffer were from BioRad (Hercules, CA). Human heparin cofactor II (HCII), antithrombin (AT), prothrombin (FII), thrombin (FIIa), factor VIIa (FVIIa), factor IX (FIX), factor IXa (FIXa), factor X (FX), factor Xa (FXa), factor XI (FXI), factor XIa (FXIa), factor XIII (FXIII), factor XIIIa (FXIIIa), and plasmin (PL) were obtained from Haematologic Technologies (Essex Junction, VT). Pooled normal human plasma was from George King Biomedical Inc (Overland Park, Kansas). Human serotransferrin, alpha-2-antiplasmin, fibronectin, and clusterin were from SinoBiological (Wayne, PA). *N*-acetylmuramoyl-L-alanine amidase was from Cusabio (Houston, TX). RED-tris-NTA $2^{nd}$ generation His-Tag labeling kit and premium capillary chips for microscale thermophoresis (MST) experiments were from NanoTemper Technologies GmbH (Munich, Germany).

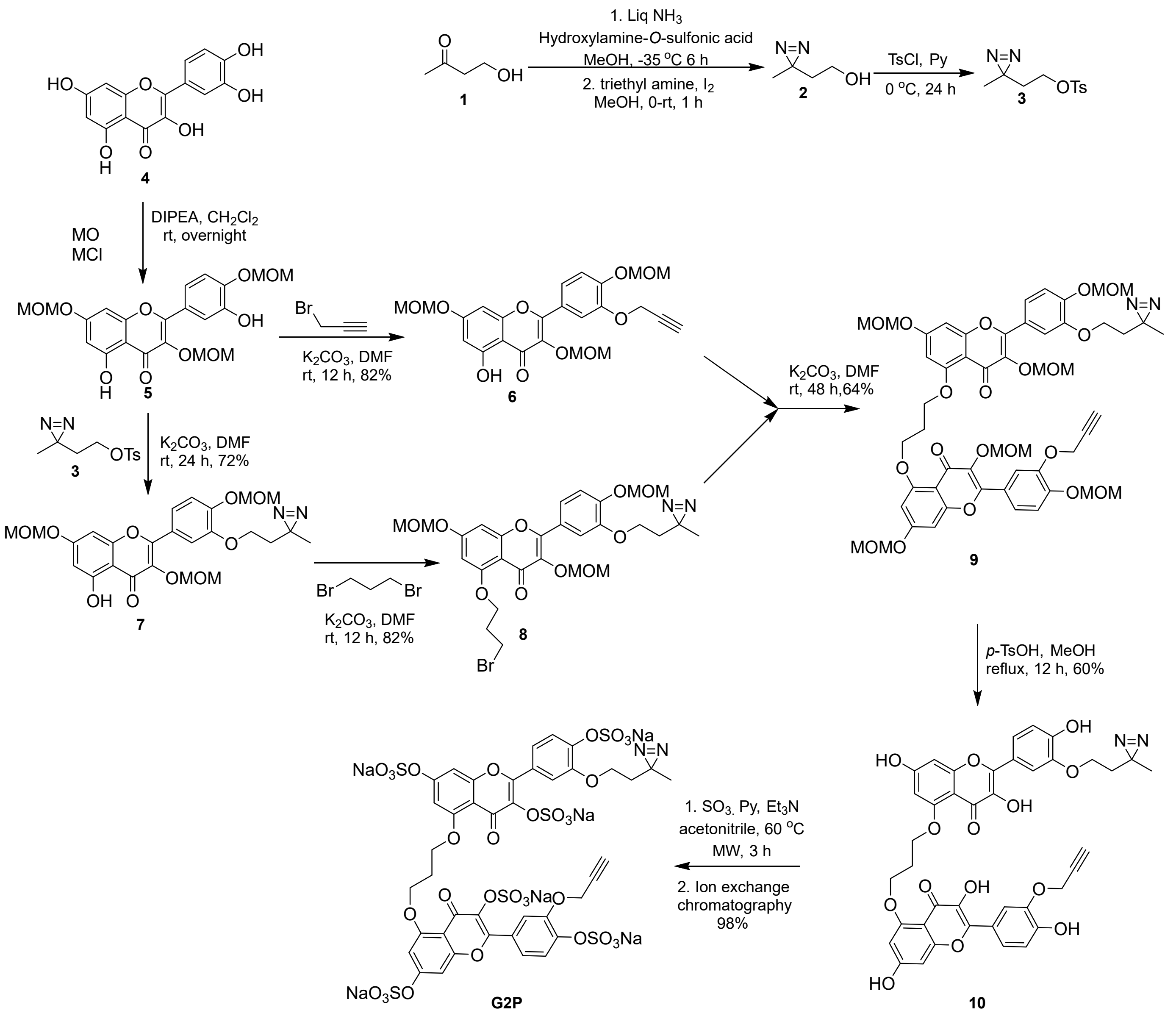


**Scheme 1. Synthesis of G2P.**

### Experimental Methods

**Synthesis**—Chemical reactions sensitive to air or moisture were carried out under nitrogen atmosphere in oven-dried glassware. Reagent solutions, unless otherwise noted, were handled under a nitrogen atmosphere using syringe techniques. Flash chromatography was performed using Teledyne ISCO (Lincoln, NE). Combiflash RF was performed using disposable normal silica cartridges of 30–50 µ particle size, 230–400 mesh size and 60 Å pore size. The flow rate of the mobile phase was in the range of 18 to 35 mL/min and mobile phase gradients of ethyl acetate/hexanes and $CH_2Cl_2/CH_3OH$ were used to elute compounds.

**Compound 3** was synthesized as previously reported (Yestrepsky, B. D.; Kretz, C. A.; Xu, Y.; Holmes, A.; Sun, H.; Ginsburg, D.; Larsen, S. D.; Development of tag-free photoprobes for studies aimed at identifying the target of novel Group A Streptococcus antivirulence agents. *Bioorg. Med. Chem. Lett.* **2014**, *24*, 1538–1544).

**Synthesis of 5**—To a solution of quercetin (**4**, 1.0 eqv.) in DCM, *N*,*N*-diisopropylethylamine (8.0 eqv.) and MOM chloride (3.5 eqv.) were added under nitrogen. After vigorous stirring at 0 °C for 1 h, the reaction mixture was allowed to warm to room temperature over 2 h followed by continued stirring for 12 h. The mixture was then diluted with water (100 mL), extracted with EtOAc (200 mL), and then the organic layer was washed with water (100 mL) and dried over $NaSO_4$. The residue obtained after removal of the solvent was purified by flash column chromatography to afford tri-protected ether **5** (50% yield).

**Compound 3** was synthesized as previously reported (). Karuturi, R.; Al-Horani, R. A.; Mehta, S. C.; Gaillani, D.; Desai, U. R. Discovery of allosteric modulators of factor XIa by targeting hydrophobic domains adjacent to Its heparin-binding site. *J. Med. Chem.* **2013**, *56*, 2415–2428.

**Synthesis of 7**—To a solution of **6** (700 mg, 1.6 mmol) in dry *N,N'*-DMF (15 mL), was added $K_2CO_3$ (333 mg, 2.4 mmol) and stirred for 2 min followed by the addition of **3** (614 mg, 2.4 mmol). After stirring for 24 h at room temperature, the reaction mixture was diluted with EtOAc (25 mL) and water (25 mL). The organic layer was separated, and the aqueous phase extracted with EtOAc (2 × 25 mL). The combined organic extracts were washed with saturated NaCl solution (25 mL) and dried over anhydrous $Na_2SO_4$. Removal of the solvent under reduced pressure followed by flash column chromatography yielded **7** in 72% yield. $^1$H NMR (400 MHz, $CDCl_3$): $\delta$ = 12.4 (s, 1H), 7.06-7.57 (m, 2H), 7.16 (d, $J$ = 8.3 Hz, 1H), 6.55 (d, $J$ = 2.1 Hz, 1H), 6.39 (d, $J$ = 2.1 Hz, 1H), 5.24 (s, 2H), 5.16 (s, 2H), 5.11 (s, 2H), 3.95 (t, $J$ = 6.5 Hz, 2H), 3.47 (s, 3H), 3.42 (s, 3H), 3.16 (s, 3H), 1.82 (t, $J$ = 6.4 Hz, 2H), 1.09 (s, 3H) ppm. $^{13}$C NMR (100 MHz, $CDCl_3$): $\delta$ = 177.6, 162.0, 160.9, 155.7, 155.5, 148.3, 147.5, 134.6, 123.5, 122.0, 115.3, 113.5, 105.6, 98.8, 96.9, 94.3, 93.2, 93.1, 63.4, 56.8, 55.4, 55.3, 33.4, 23.1, 19.2 ppm. HR-MS (Q-ToF): *m/z*: $[M + H]^+$ calculated for $C_{25}H_{28}O_{10}N_2$ 516.1743, found 517.1778.

**Synthesis of 11**—To a solution of **7** (620 mg, 1.20 mmol) in dry *N,N'*-DMF (15 mL), was added $K_2CO_3$ (830 mg, 6.00 mmol) and the mixture stirred for 2 min followed by the addition of 1,3-dibromopropane (1.21 g, 6.00 mmol). After stirring for 12 h at room temperature, the reaction mixture was diluted with EtOAc (25 mL) and water (25 mL). The organic layer was separated, and the aqueous phase extracted with EtOAc (2×25 mL). The combined organic extracts were washed with saturated NaCl solution (25 mL) and dried over anhydrous $Na_2SO_4$. Removal of the solvent under reduced pressure followed by flash column chromatography yielded **11** in 97% yield. $^1$H NMR (400 MHz, $CDCl_3$): $\delta$ = 7.59 (d, $J$ = 2.0 Hz, 1H), 7.56 (dd, $J$ = 8.5, 1.9 Hz, 1H), 7.14 (d, $J$ = 8.5 Hz, 1H), 6.65 (d, $J$ = 2.1 Hz, 1H), 6.39 (d, $J$ = 2.2 Hz, 1H), 5.22 (t, $J$ = 2.2 Hz, 2H), 5.18 (s, 2H), 5.11 (s, 2H), 4.14 (t, $J$ = 5.6 Hz, 2H), 3.96 (t, $J$ = 5.9 Hz, 2H), 3.75 (t, $J$ = 5.6 Hz, 2H), 3.47 (s, 3H), 3.44 (s, 3H), 3.13 (s, 3H), 2.26 (t, $J$ = 6.00 Hz, 2H), 1.81 (t, $J$ = 6.5 Hz, 2H), 1.08 (s, 3H) ppm. $^{13}$C NMR (100 MHz, $CDCl_3$): $\delta$ = 172.5, 160.4, 159.2, 157.5, 152.5, 147.7, 147.4, 136.9,

124.0, 121.6, 115.4, 113.5, 109.2, 96.9, 96.8, 94.6, 94.4, 93.3, 65.6, 63.3, 56.6, 55.4, 55.3, 33.4, 30.9, 29.6, 23.1, 19.2 ppm. HR-MS (Q-ToF): *m/z*: $[M+Na]^+$ calculated for $C_{28}H_{33}BrN_2O_{10}Na$ 659.1216, found 659.1204.

**Synthesis of 13**—To a solution of **11** (350 mg, 0.74 mmol) in dry *N,N'*-DMF (15 mL), was added $K_2CO_3$ (307 mg, 2.22 mmol) and the mixture stirred for 2 min followed by the addition of **12** (614 mg, 0.93 mmol). After stirring for 48 h at room temperature, the reaction mixture was diluted with EtOAc (25 mL) and water (25 mL). The organic layer was separated, and the aqueous phase was extracted with EtOAc (2×25 mL). The combined organic extracts were washed with saturated NaCl solution (25 mL) and dried over anhydrous $Na_2SO_4$. Removal of the solvent under reduced pressure followed by flash column chromatography yielded **13** in 64% yield. $^1$H NMR (400 MHz, $CDCl_3$): $\delta$ = 7.75 (d, *J* = 2.1 Hz, 1H), 7.60-7.53 (m, 3H), 7.17 (d, *J* = 8.7 Hz, 1H), 7.13 (d, *J* = 8.6 Hz, 1H), 6.57 (t, *J* = 2.4 Hz, 2H), 6.47 (d, *J* = 1.7 Hz, 2H), 5.2 (s, 4H), 5.14 (d, *J* = 1.1 Hz, 4H), 5.11 (d, *J* = 3.1 Hz, 4H), 4.76 (d, *J* = 2.5 Hz, 2H), 4.38 (t, *J* = 5.4 Hz, 4H), 3.94 (t, *J* = 6.4 Hz, 2H), 3.46 (d, *J* = 2.5 Hz, 6H), 3.40 (s, 6H), 3.12 (d, *J* = 1.2 Hz, 6H), 2.46-2.43 (m, 3H), 1.80 (t, *J* = 6.4 Hz, 2H), 1.08 (s, 3H) ppm. $^{13}$C NMR (100 MHz, $CDCl_3$): $\delta$ = 173.6, 161.5, 160.5, 158.4, 153.2, 153.1, 149.0, 148.6, 148.4, 147.1, 137.9, 137.8, 125.1, 124.9, 123.4, 122.6, 116.4, 115.9, 115.7, 114.5, 110.1, 98.0, 97.9, 97.8, 95.4, 95.3, 95.2, 95.1, 94.3, 94.2, 78.3, 77.3, 75.9, 65.7, 64.3, 57.6, 57.5, 57.1, 56.4, 56.3, 34.4, 28.9, 24.1, 20.1 ppm. HR-MS (Q-ToF): *m/z*: $[M + Na]^+$ calculated for $C_{52}H_{56}N_2O_{20}Na$ 1051.3324, found 1051.3309.

**Synthesis of 14**—To a solution of **13** (400 mg, 0.38 mmol) in MeOH (20 mL) was added *p*-toluenesulfonic acid (1.7 g, 7.78 mmol) at room temperature under $N_2$ atmosphere and the mixture stirred for 12 h at reflux. At the end of the reaction (TLC monitoring), the reaction mixture was filtered, washed with excess EtOAc and dried to obtain polyphenol **14** as yellow solid in 50% yield. $^1$H NMR (400 MHz, DMSO): $\delta$ = 10.6 (s, 2H), 9.76 (s, 1H), 9.58 (s, 1H), 8.76 (s, 2H), 7.81 (d, *J* = 2.1 Hz, 1H), 7.73 (dd, *J* = 9.6, 2.0 Hz, 2H), 7.66 (dd, *J* = 8.7, 2.0 Hz, 1H), 6.96 (d, *J* = 8.4 Hz, 2H), 6.52 (t, *J* = 2.2 Hz, 2H), 6.39 (d, *J* = 1.5 Hz, 2H), 4.85 (d, *J* = 2.2 Hz, 2H), 4.37 (t, *J* = 6.0 Hz, 4H), 3.96 (t, *J* = 6.4 Hz, 2H), 3.56 (t, *J* = 2.5 Hz, 1H), 2.32 (t, *J* = 5.6 Hz, 2H), 1.82 (t, *J* = 6.2 Hz, 2H), 1.13 (s, 3H) ppm. $^{13}$C NMR (100 MHz, DMSO): $\delta$ = 171.0, 162.4, 159.7, 157.8, 148.7, 148.6, 146.2, 145.1, 141.6, 141.4, 137.3, 137.2, 122.3, 122.2, 122.1, 121.4, 116.0, 115.8, 114.2, 113.4, 105.2, 96.4, 94.7, 79.3, 78.3, 64.8, 64.1, 56.5, 33.7, 24.5, 19.7 ppm. HR-MS (Q-ToF): *m/z*: $[M + K]^+$ calculated for $C_{40}H_{32}N_2O_{14}Na$ 787.1751, found 787.1709.

**Synthesis of 15**—To a stirring solution of **14** (40 mg, 0.052 mmol) in anhydrous $CH_3CN$ (3 mL) at room temperature, $Et_3N$ (0.317 mL, 3.14 mmol) and $SO_3$/pyridine complex (299 mg, 1.88 mmol) were added. The reaction vessel was sealed and microwaved (CEM Discover, Cary, NC) for 4 h at 60 ºC followed by cooling and concentration *in vacuo* at <30 °C. The mixture was then purified on Combiflash RF system using 5-15% $CH_3OH$ in $CH_2Cl_2$ mobile system to obtain the per-sulfated compound in quaternary ammonium salt form. Fractions containing the desired molecule (per-sulfation) were pooled, concentrated *in vacuo*, and re-loaded onto a SP Sephadex C-25 column for sodium exchange. Fractions containing the per-sulfated molecule in sodiated form were pooled and lyophilized to obtain a fluffy off-white powder **15** in 97% yield. $^1$H NMR (400 MHz, DMSO): $\delta$ = 8.00 (dd, *J* = 6.9, 2.0 Hz, 2H), 7.68 (dd, *J* = 8.8, 2.0 Hz, 1H), 7.62 (dd, *J* = 8.7, 1.9 Hz, 1H), 7.57 (dd, *J* = 8.7, 2.4 Hz, 2H), 7.07 (t, *J* = 2.0 Hz, 2H), 6.71 (dd, *J* = 5.6, 1.9 Hz, 2H), 4.78 (d, *J* = 2.2 Hz, 2H), 4.24 (t, *J* = 6.4 Hz, 4H), 3.99 (t, *J* = 6.5 Hz, 2H), 3.47 (t, *J* = 2.3 Hz, 1H), 2.35-2.24 (m, 2H), 1.67 (t, *J* = 6.3 Hz, 2H), 1.07 (s, 3H) ppm. $^{13}$C NMR (100 MHz, DMSO): $\delta$ = 171.5, 162.9, 160.2, 158.3, 149.2, 149.0, 146.7, 145.6, 142.0, 141.9, 137.8, 137.7, 122.7, 122.6, 122.5, 121.8, 116.4, 116.2, 114.5, 113.7, 105.6, 99.7, 95.2, 79.8, 78.9, 65.2, 64.4, 56.9, 34.1, 29.0, 25.1, 20.2 ppm. MS (ESI) calculated for $C_{40}H_{26}N_2Na_6O_{32}S_6$ $[(M-2Na)/2]^{2-}$, m/z 664.9197, found for $[(M-2Na)/2]^{2-}$, m/z 664.8884.

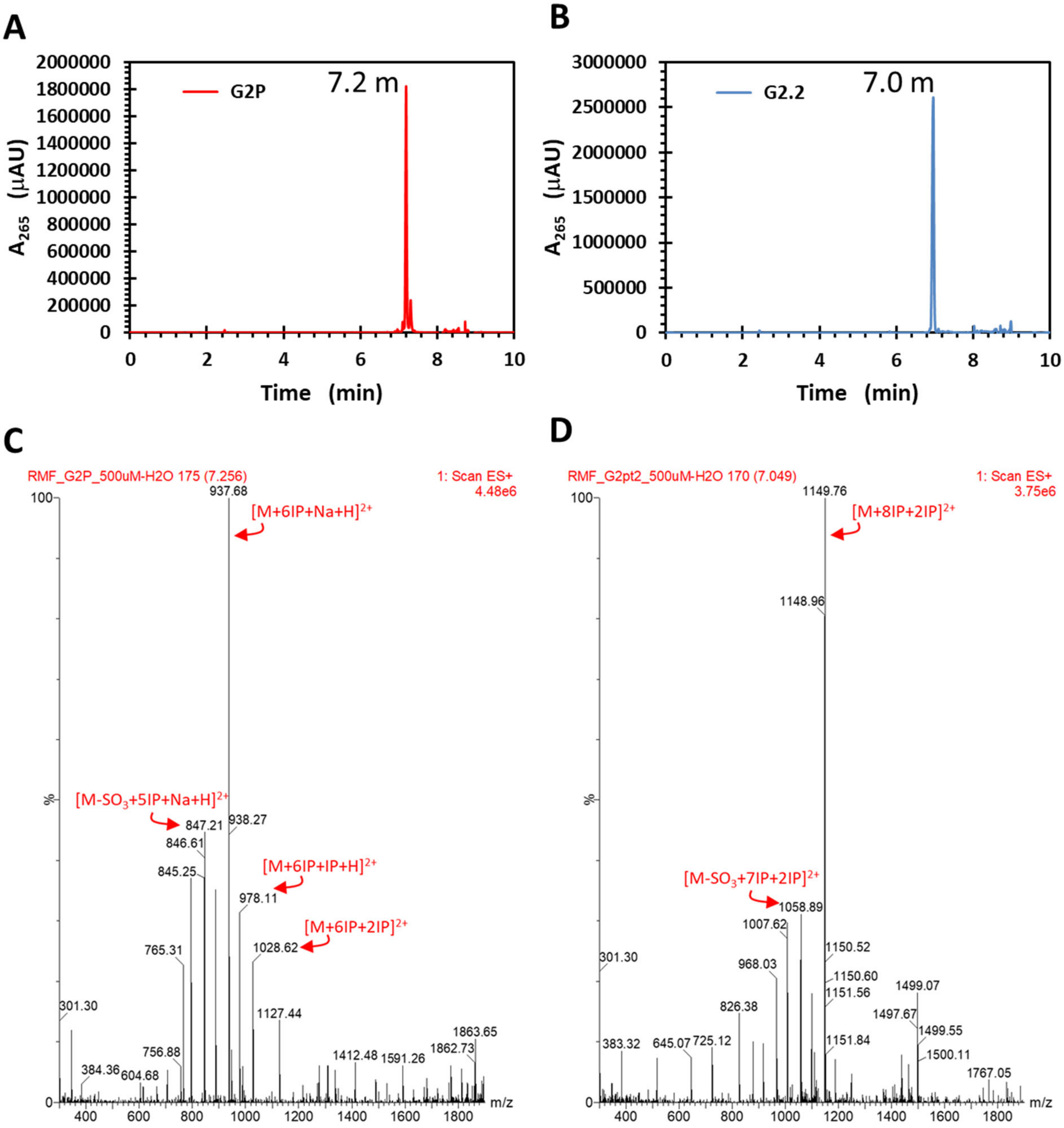


**Figure S1**. (A & B) Reversed-phased ion-pairing (RP-IP) ultrahigh pressure liquid chromatography (UPLC) and (C & D) mass spectrometry (MS) profile of G2P and G2.2, respectively. Profiles A show the purity of G2P and G2.2 to be >95%, whereas the mass peaks confirm their structures, which includes 6 sulfate groups, a diazirine and an alkyne for G2P and 8 sulfate groups for G2.2. IP refers to ion-pairing agent, which is hexylamine ($CH_3(CH_2)_5NH_3^+$).

**Photoaffinity labeling of proteins in buffer**—For photolabeling of individual proteins, G2P was added to a solution of the target protein (~0.2 μM) in buffer so as to yield 0—100 μM G2P concentrations, as shown in **Tables S1**. For photolabeling of a mixture of proteins, 30 μg each protein was first mixed in an Eppendorf tube and the solution volume made up to 100 μL with PBS. Ten μL volume of this mixture was diluted to 200 μL with PBS and **G2P** was added to achieve the desired concentration (0–100 μM), as shown in **Table S2**. The procedure to analyze individual proteins or a mixture of proteins was identical. The samples were mixed gently with a pipette and incubated in the dark at room temperature for 30 min, followed by UV irradiation at 368 nm for 30 min at 4 ºC. AF488 (1.5×[G2P] μM), TBTA (1.5×[G2P] μM), TCEP (15×[G2P] μM), and $CuSO_4$ (15×[G2P] μM) was then added and the solution incubated at room temperature for 2 h. To each sample was then added 1.2 mL of pre-chilled acetone (-20 ºC), followed by gentle vortexing and overnight incubation at -80 ºC. Centrifugation at 17,000g and 4 ºC for 15 min yielded a protein pellet, which was washed twice with cold methanol and then used for SDS-PAGE analysis per standard protocol. The SDS-PAGE gel was washed thoroughly with $ddH_2O$ to remove unreacted AF488 and imaged using a GE gel imager. Coomassie or silver staining was performed followed by white light or fluorescence imaging. Band intensities were determined using Quantity One software from Bio-Rad.

**Table S1.** Concentrations used in G2P-labeling of FIIa and FIXa.

| | **FIIa (thrombin)** | | | | | **FIXa** | | | | |
|---|---|---|---|---|---|---|---|---|---|---|
| | **1** | **2** | **3** | **4** | **5** | **1** | **2** | **3** | **4** | **5** |
| **[Protein]# (μM)** | 2 | 2 | 2 | 2 | 2 | 2 | 2 | 2 | 2 | 2 |
| **[G2.2-P] (μM)** | 0 | 0.5 | 2 | 20 | 100 | 0 | 0.5 | 2 | 20 | 100 |

**Table S2.** Conditions for G2P-labeling of protein mix.

| **Conditions** | **1** | **2** | **3** | **4** | **5** | **6** | **7** | **8** |
|---|---|---|---|---|---|---|---|---|
| **Protein Mix# (μL)** | 10 | 10 | 10 | 10 | 10 | 10 | 10 | 10 |
| **PBS buffer (μL)** | 190 | 190 | 190 | 190 | 190 | 190 | 190 | 190 |
| **[G2.2] (μM)** | 0 | 0 | 0 | 0 | 0 | 0 | 300 | 0 |
| **[G2P] (μM)** | 0 | 1 | 5 | 10 | 25 | 25 | 10 | 0 |
| **UV irradiation&** | + | + | + | + | + | - | + | + |
| **[AF488] (μM)** | 37.5 | 1.5 | 7.5 | 15 | 37.5 | 37.5 | 15 | 0 |

#Protein mix consisted of human hemoglobin, heparin cofactor II, prothrombin, antithrombin III, BSA, IGF-1, and factors IX, X, XI, and XIII (~3 mg each); &UV irradiation = exposure of sample to 368 nm light for 30 min at 4 °C.

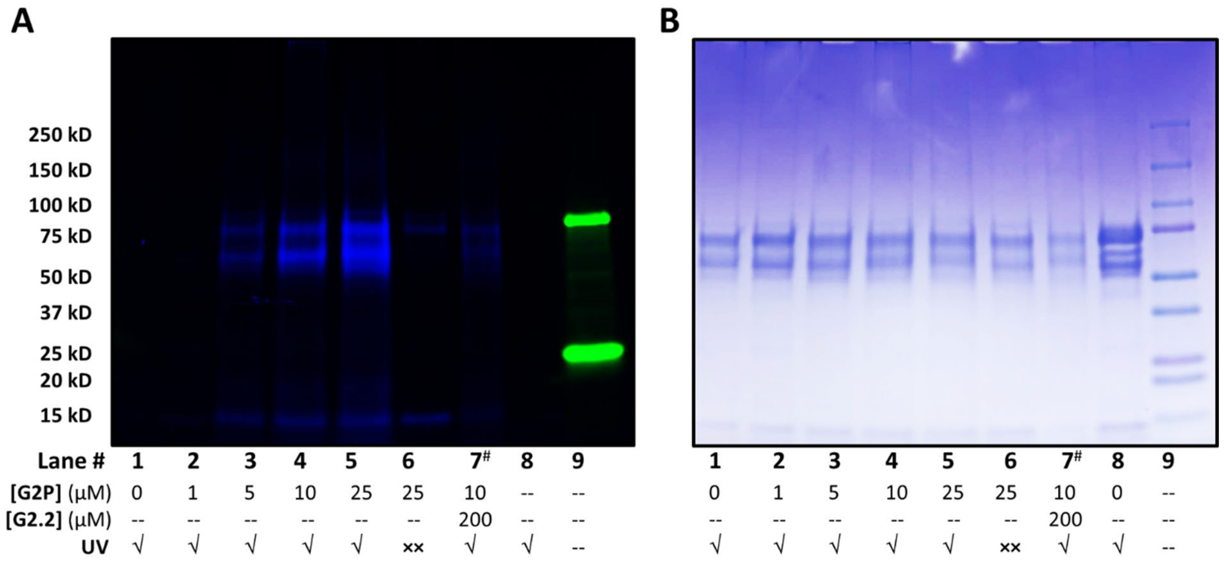


**Figure S2**. PAGE analysis of rhodamine(Rh)-coupled products following photolysis ($\lambda_{EX}$ = 368 nm) of a mixture of 10 proteins (3 μg each) and G2P (0→25 μM) for 10 min followed by reaction with Rh-$N_3$ (1.5×[G2P] μM). Lanes 1 to 9 contain samples with different levels of G2P (0→100 μM) treated with or without irradiation (shown as either √ or ××, respectively) in the presence (Lane 7) or absence of G2.2 (200 μM) as a competitor of G2P. The protein mix contained IGF1, factor IX, antithrombin, factor X, hemoglobin, heparin cofactor II, BSA, prothrombin, factor XI, and factor XIII. (A) Fluorescence image using blue and green filters was saved for off-line image analysis using Image J. (B) Coomassie-stained gel image.

**Depletion of Abundant Plasma Proteins**—Depletion of proteins present in human plasma in abundance was performed using Mini Top-12 Abundant Protein Depletion Spin Columns according to the manufacturer's recommendations (ThermoFisher). Briefly, human plasma (10 µL) was applied to each column and incubated at room temperature for 30 min with gentle end-over-end mixing. Plasma depleted in abundant proteins was obtained as the flow-through by centrifugation at 1000g for 2 min.

**G2P photolabeling of human plasma proteins**—Ten µL of human plasma, depleted of abundant proteins, was added to a solution of G2P to achieve 0—75 µM concentration of the probe, as shown in **Table S3**. The samples were mixed gently with a pipette and incubated in the dark at room temperature for 30 min, followed by UV irradiation at 368 nm for 30 min at 4 °C. AF488 (1.5×[G2P] µM), TBTA (1.5×[G2P] µM), TCEP (15×[G2P] µM), and $CuSO_4$ (15×[G2P] µM) was then added and the solution incubated at room temperature for 2 h. To each sample was then added 1.2 mL of pre-chilled acetone (-20 °C), followed by gentle vortexing and overnight incubation at -80 °C. Centrifugation at 17,000g and 4 °C for 15 min yielded a protein pellet, which was washed twice with cold methanol and then used for SDS-PAGE per standard protocol. The gel was washed thoroughly with $ddH_2O$ to remove unreacted AF488 and imaged using a GE gel imager. Coomassie or silver staining was performed after fluorescence imaging.

**Table S3.** Conditions for G2P-labeling of human plasma proteins.

| | **1** | **2** | **3** | **4** | **5** | **6** | **7** | **8** | **9** |
|---|---|---|---|---|---|---|---|---|---|
| **Human plasma[#] (µL)** | 10 | 10 | 10 | 10 | 10 | 10 | 10 | 10 | 10 |
| **Protein depleted plasma[$]** | + | + | + | + | + | + | + | + | - |
| **[G2.2] (µM)** | 0 | 0 | 0 | 0 | 0 | 0 | 1500 | 0 | 0 |
| **[G2P] (µM)** | 0 | 10 | 25 | 50 | 100 | 100 | 50 | 0 | 0 |
| **UV irradiation[&]** | + | + | + | + | + | - | + | + | + |
| **[AF488] (µM)** | 150 | 15 | 37.5 | 75 | 150 | 150 | 75 | 0 | 0 |

[#]Human plasma consisted of pooled plasma from George King Biomedical; [$]Pooled human plasma was depleted of its most abundant protein using spin columns (ThermoFisher Scientific, Catalog No: 85165); [&]UV irradiation = exposure of sample to 368 nm light for 30 min at 4 °C.

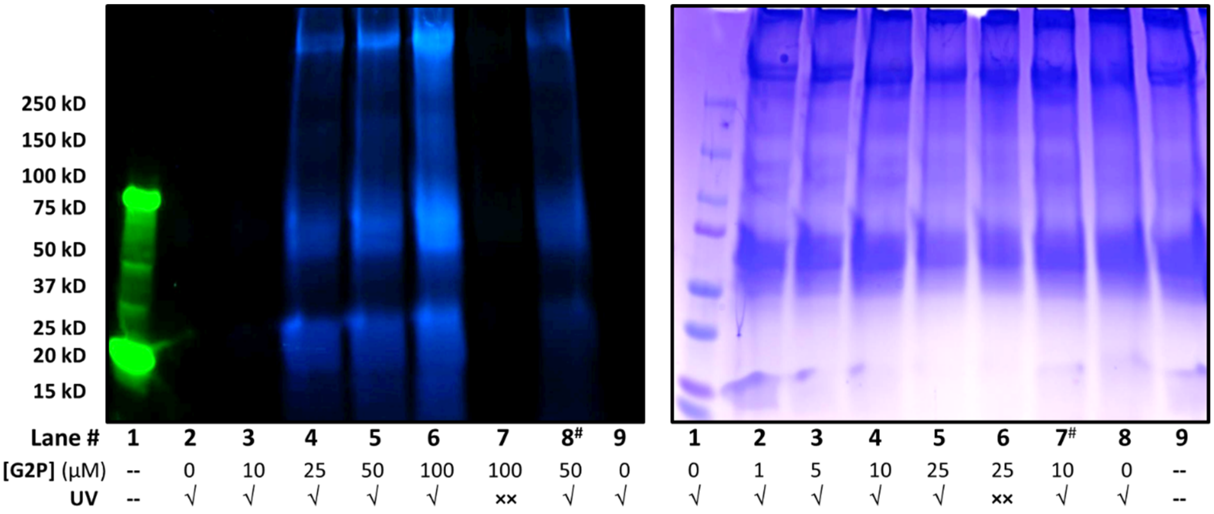


**Figure S3.** Fluoresence image of SDS-PAGE gel of rhodamine(Rh)-coupled products following photolysis ($\lambda_{EX}$ = 368 nm) of pooled human plasma samples containing G2P followed by coupling with Rh-$N_3$. Lanes 1 to 9 contain samples with different levels of G2P (0→100 μM) treated with or without irradiation (shown as either √ or ××, respectively) in the presence of 1500 μM unlabeled G2.2 (Lane 8; marked #) or its absence. Human plasma was first passed through Pierce™ Top-12 depletion spin columns to deplete most abundant proteins (> 90%), then treated with G2P and photolyzed. (Right) Coomassie-stained gel image. (Left) Fluorescence image using blue and green filters was saved for off-line image analysis using Image J.

**Biotin affinity pull-down and SDS-PAGE of G2P-labeled human plasma proteins**— Ten µL of human plasma, depleted of abundant proteins, was added to a solution of G2P to achieve 0—75 µM concentration of the probe, as shown in **Table S4**. The samples were mixed gently with a pipette and incubated in the dark at room temperature for 30 min, followed by UV irradiation at 368 nm for 30 min at 4 ºC. AF488 (1.5×[G2P] µM), TBTA (1.5×[G2P] µM), TCEP (15×[G2P] µM), and $CuSO_4$ (15×[G2P] µM) was then added and the solution incubated at room temperature for 2 h. To each sample was then added 1.2 mL of pre-chilled acetone (-20 ºC), followed by gentle vortexing and overnight incubation at -80 ºC. Centrifugation at 17000g and 4 ºC for 15 min yielded a protein pellet, which was washed twice with cold methanol and resuspended in 100 µL PBS buffer containing 1% (w/v) SDS. Then, 400 µL PBS was added to each sample to dilute SDS to 0.2% and the samples transferred to Eppendorf tubes containing 300 µL of pre-washed high-capacity streptavidin agarose beads. Following incubation at 4 ºC with gentle end-over-end mixing for 2 h, the samples were centrifuged at 1000g for 5 min. The supernatant containing the unbound fraction was collected, while the beads were washed three times with PBS buffer (1 mL) containing 0.2% SDS (w/v) to ensure removal of unbound proteins. To the washed beads was added 250 µL of PBS containing 4% SDS followed by boiling at 95 ºC for 15 min, and centrifugation at 13000g. The supernatant containing the bound fraction was collected. The bound and unbound fractions were then subjected to SDS-PAGE analysis, as per standard protocols with silver staining. Proteomics analysis was also performed on the bound fractions.

**Table S4.** Conditions for G2P labeling and affinity pulldown of proteins from human plasma.

| | | | | | |
|---|---|---|---|---|---|
| **Human plasma (µL)** | 10 | 10 | 10 | 10 | 10 |
| **Top-2 protein depletion** | + | + | + | + | + |
| **[G2P] (µM)** | 0 | 7.5 | 15 | 30 | 75 |
| **[Azido-PEG-Biotin] (µM)** | 112.5 | 11.2 | 22.5 | 45 | 112.5 |

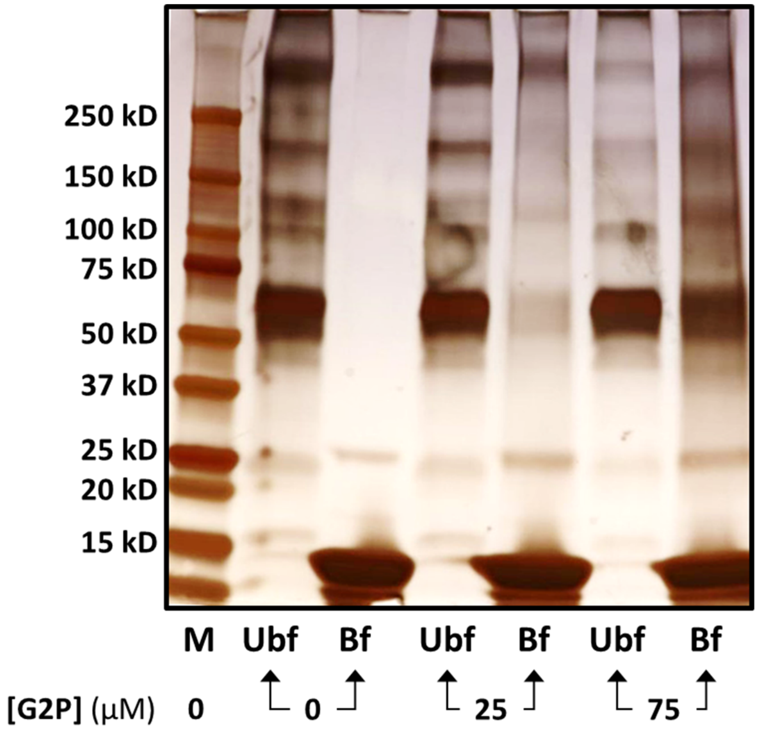


**Figure S4.** Human plasma was first passed through Pierce™ Top-12 depletion spin columns to deplete most abundant proteins (> 90%), then treated with G2P and photolyzed. Image of silver-stained SDS-PAGE gel of biotin(Bt)-coupled products following photolysis ($\lambda_{EX}$ = 368 nm) of human plasma samples containing G2P (0, 25, or 75 μM) followed by reaction with Bt-$N_3$. The Bt-coupled products were passed over a streptavidin affinity matrix to collect the unbound fraction (Ub) and then eluted with 4% SDS to collect the bound fraction (B) at each concentration of G2P.

**Proteomics Sample Preparation**—To a sample (400 µL) from biotin-affinity pull-down was added 45 mM dithiothreitol (DTT, 44.4 µL) in 100 mM triethylammonium bicarbonate (TEAB), pH 8.5, and incubated at 37 °C for 1 h. After allowing the samples to cool, 100 mM iodoacetamide (IAA, 49.4 µL) in 100 mM TEAB was added and the samples incubated at room temperature in the dark for 30 min. Unreacted IAA was quenched by adding 100 mM DTT (54.9 µL) in 100 mM TEAB. Samples were then acidified with 12% (v/v) *o*-phosphoric acid (61 µL), transferred to centrifuge tubes followed by introduction of LC-MS grade methanol (5 mL) and overnight incubation at -80°C to precipitate proteins, which pelleted by centrifugation (30 min, 4000g). All, but ~500 µL, of the supernatant liquid was removed and the proteins homogenized by vigorous vortexing and loaded onto an S-Trap micro spin column (120 µL at a time) by centrifuging at 1000g for 1 min. The column was then washed four times with LC-MS grade methanol (150 µL) with centrifugation at 1000g for 1 min each. Each spin column was then treated with MS grade trypsin (2 µg in 25 µL 50 mM TEAB) and incubated overnight at 37 °C. A second dose of trypsin (2 µg in 25 µL 50 mM TEAB) was introduced into onto each spin column and incubated at 37 °C for 2 h. Peptides from trypsin digestion were recovered by sequential washing of the spin column with 25 µL of 50 mM TEAB, then 25 µL of solvent A (2:98 LC-MS grade acetonitrile: LC-MS grade water supplemented with 0.1% (v/v) formic acid), and 25 µL of solvent B (80:20 LC-MS grade acetonitrile: LC-MS grade water supplemented with 0.1% (v/v) formic acid). Acetonitrile was removed using a centrifugal vacuum concentrator and peptide concentration was measured through absorbance at 215 nm. Samples were diluted to 0.5 g/L using solvent A and 2 µL of the solution (1 µg total protein) was used for LC-MS/MS analysis.

**Proteomics LC-MS Experimentation**—Samples were loaded onto a pre-column Acclaim PepMap 100 (Thermo Scientific, 100 µm × 2 cm), which was followed by an analytical column (50 cm µPAC (PharmaFluidics). A flow rate of 150 nL/min was set up using an Easy-nLC 1200 (Thermo Scientific) UPLC/autosampler. Peptides were eluted utilizing a 2 to 45% gradient of solvent B (80:20 LC-MS grade acetonitrile: LC-MS grade water supplemented with 0.1% (v/v) formic acid) in solvent A (2:98 LC-MS grade acetonitrile: LC-MS grade water supplemented with 0.1% (v/v) formic acid) over 44 min. Following set up was used for peptide analysis: Spray voltage on the µPAC compatible Easy-Spray emitter was set to 1300 volts; ion transfer tube temperature was fixed at 275 °C; RF lens was set to 30%; and the default charge state was set to 3. MS data in the range of 400—1500 m/z were collected using the orbitrap at 120000 resolution in positive mode with an AGC target of $4.0\times10^5$ and a maximum injection time of 50 ms. Peaks were filtered for MS/MS analysis based on an isotopic peak distribution expected for a known peptide with an intensity above $2.0\times10^4$ and a charge state of 2 to 5. Peaks were excluded dynamically with the MS/MS set to be collected at 45% of chromatographic peak width. MS/MS peaks of >150 m/z were collected using the Orbitrap at 15000 resolution in positive centroid mode with an AGC target of $1.0\times10^5$ and a maximum injection time of 200 ms. Activation type was HCD stepped from 29 to 31. The MS and MS/MS data were analyzed utilizing several different software packages including Proteome Discoverer 2.4, Mascot Server 2.7 and Byonic. Initially, a simple search to identify proteins present in the sample was performed using Proteome Discoverer combining a Sequest HT and Mascot search into one summative result for each sample. Both searches utilized the UniProt human proteome reference database (downloaded May, 11 2020) and a common protein contaminant database available within Proteome Discoverer. Each search assumed trypsin-specific peptides with the possibility to 2 missed cleavages, a precursor mass tolerance of 10 ppm and a fragment mass tolerance of 0.1 Da. Both searches included common static peptide residue modifications including carbamidomethylation of Cys, oxidized Met, acetylated *N*-termini, and cyclized *N*-terminal Gln to pyro-Glu. The Sequest HT search also included dynamic modifications at the *N*-terminus of a protein due to the loss of Met, which may be followed by *N*-terminal acetylation. The proteins obtained from these searches (available as Supplementary Materials; files 201022_VCU_A_1.xlsx and 201022_VCU_B_1.xlsx) were further analyzed using Byonic. In this analysis, static or dynamic modifications previously not known were sought to be identified. This relied on searching for any mass shift(s) of 10—2000 from those identified in trypsin-specific peptide analysis performed in the first search (see two Excel spreadsheets containing proteomics data, which can be downloaded from the publisher's website). For these to be reliable, analytic parameters including the score and delta score have to be high (>300), which should be followed by manual analysis of each MS/MS spectra for high-fidelity validation.

**Proteomics identification of high-fidelity G2P targets**—To exclude false positives in the set of proteins identified by the above proteomics protocol, the following stringent criteria were applied to every experimental conditions: 1) Proteins identified had to pass a high combined protein false discovery rate (FDR) confidence interval; 2) Proteins had to have a combined experimental q-value of 0; 3) Proteins had to have a minimum of two or more peptide spectrum matches; 4) Proteins had to have SEQUEST and Mascot scores of >0 and >30, respectively; and 5) Proteins had to have at least one unique peptide detected in proteomics experiments. Finally, all proteins identified in the control experiment ([G2P] = 0 μM) were excluded from the proteins identified at each of the test concentrations of G2P, i.e., 7.5, 15, 30, and 75 μM. Proteins common between different experimental conditions were identified using Venny (Oliveros, J. C. (2007-2015) Venny. An interactive tool for comparing lists with Venn's diagrams. https://bioinfogp.cnb.csic.es/tools/venny/index.html).

**Search for GAG-binding consensus sequences**—The sequences of identified proteins were downloaded from the UniProt database into MS word, and Lys (K), Arg (R), and His (H) residues were color coded (red). Cardin-Weintraub GAG-binding consensus sequences (Cardin AD, Weintraub HJ. Molecular modeling of protein-glycosaminoglycan interactions. Arteriosclerosis. 1989 Jan-Feb;9(1):21-32) for the proteins were identified by conducting a manual inspection of each protein sequence.

**Binding affinity of His-tagged protein for RED-tris-NTA 2nd generation dye**—The affinity of a His-tagged protein for RED-tris-NTA 2nd generation dye was first measured. Briefly, sufficient volume of 4 μM protein in PBST was first prepared. Then a 16-step, 2-fold serial dilution protocol was employed in a 384-well plate to prepare 10 μL samples of each dilution added to 10 μL of 50 nM dye in PBST. The samples were mixed, incubated in dark at room temperature for 30 min, and then loaded into Monolith NT.115 capillaries for microscale thermophoresis (MST) experiments. MST experiments were performed at 40% LED/excitation and medium MST power. The $K_D$ of the protein – dye complex was calculated using the manufacturer's in-house analytical tool (MO.Affinity Analysis).

**MST-based affinity of protein–G2.2 or protein–G2P complexes**—First, His-tagged proteins were labelled with RED-tris-NTA 2nd generation dye according to the manufacturer's instructions. For a protein with a high dye affinity (i.e., $K_D$ <10 nM), 100 μL of protein (200 nM) in PBST was mixed with 100 μL of 100 nM RED-tris-NTA 2nd generation dye in PBST. For a protein with moderate dye affinity (i.e., $K_D$ >10 nM), 100 μL of the protein (20×$K_D$ concentration) was mixed with 100 μL of 100 nM RED-tris-NTA 2nd generation dye. The sample was mixed gently and incubated at room temperature in the dark for 30 min followed by centrifugation at 15000g for 10 min. The labelled protein was transferred to a fresh Eppendorf tube. A 12-step, 2-fold serial dilution of either G2.2 or G2P in PBST was prepared in a 384-well plate with a final volume of 10 μL in each well. Then, 10 μL of a labelled protein was added to each well, mixed by gentle pipetting, incubated at room temperature in the dark for 20 min, and loaded into Monolith NT. premium capillary chips. MST experiments were carried out at 40% LED/excitation and medium MST power. The $K_D$ of the protein – dye complex was calucated using the manufacturer's in-house analytical tool (MO.Affinity Analysis).

**Table S5.** GAG-binding properties of 12 preferred protein targets based on GAG database[#] and Cardin-Weintraub (CW) primary sequence.[$]

| ID | Protein | Entry in GAG Database? | CW Sequence Present? |
|---|---|---|---|
| P00747 | Plasminogen | Y | Y |
| P02679 | Fibrinogen gamma chain | Y | Y |
| P02747 | Complement C1q subcomponent | Y | -- |
| P02751 | Fibronectin | Y | Y |
| P02787 | Serotransferrin | -- | Y |
| P04003 | C4b-binding protein alpha chain | Y | Y |
| P04275 | von Willebrand factor | Y | Y |
| P07996 | Thrombospondin-1 | Y | Y |
| P08697 | Alpha-2-antiplasmin | Y | -- |
| P10909 | Clusterin | Y | Y |
| P20851 | C4b-binding protein beta chain | Y | Y |
| Q96PD5 | *N*-acetylmuramoyl-*L*-alanine amidase | -- | Y |

[#] GAG database is described in Vallet et al. Am J Physiol Cell Physiol. 2022 Jun 1; 322(6): C1271-C1278.
[$] Cardin-Weintraub primary sequences include 'XBBXBX' and 'XBBBXXBX' sequences (X = hydropathic residue; B = Basic residue) (see Sankaranarayanan et al. Curr Opin Struct Biol. 2018 Jun; 50: 91-100).

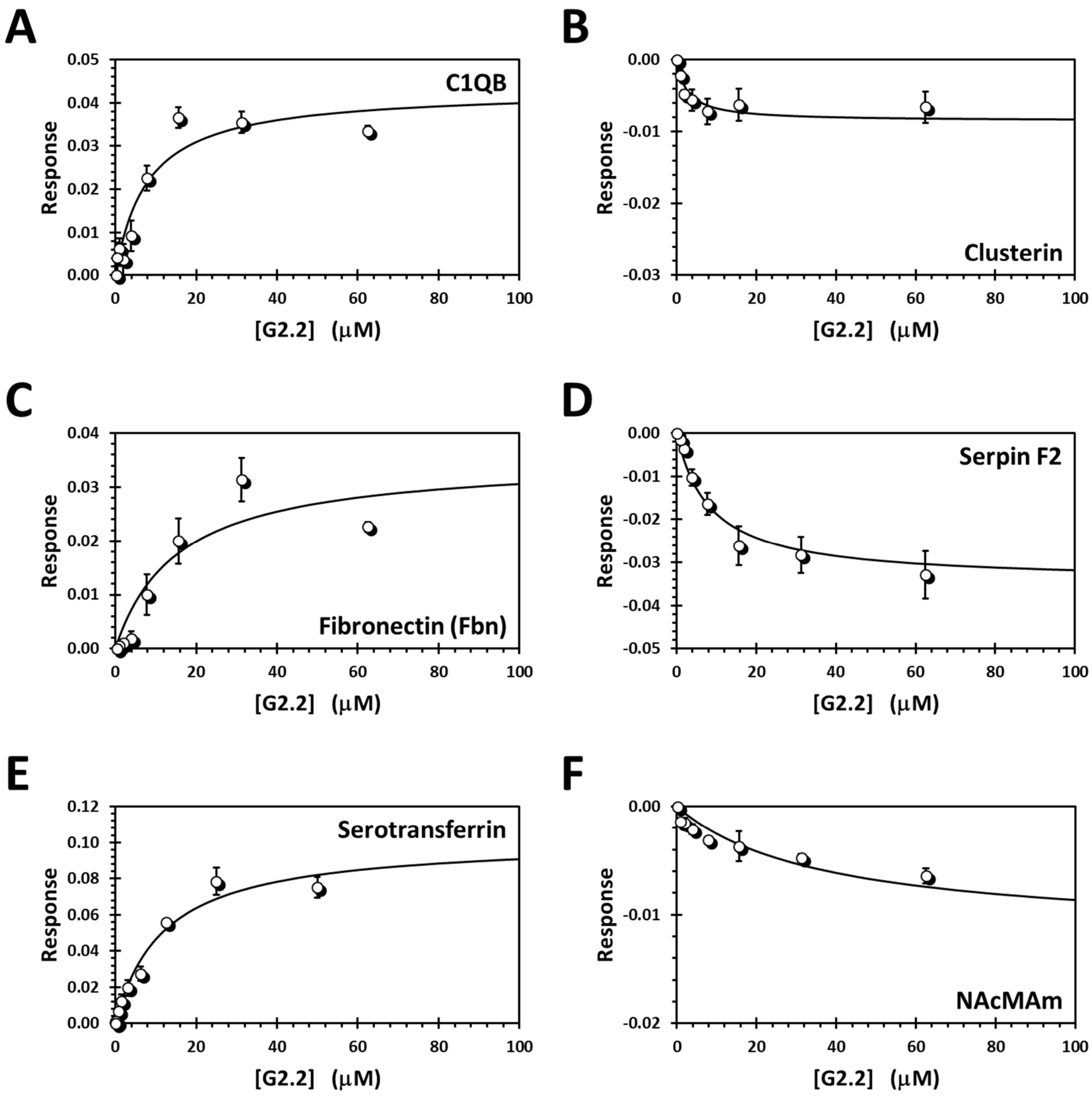


**Figure S5.** Binding affinity of G2P for the six representative protein targets identified through PAL-Pomics. His-tagged proteins, labeled with RED-tris-NTA dye, were used in microscale thermophoresis (MST)-based titrations to measure $K_D$ of interaction. Solid line shows non-linear regression using the standard hyperbolic binding equation to calculate the $K_D$.

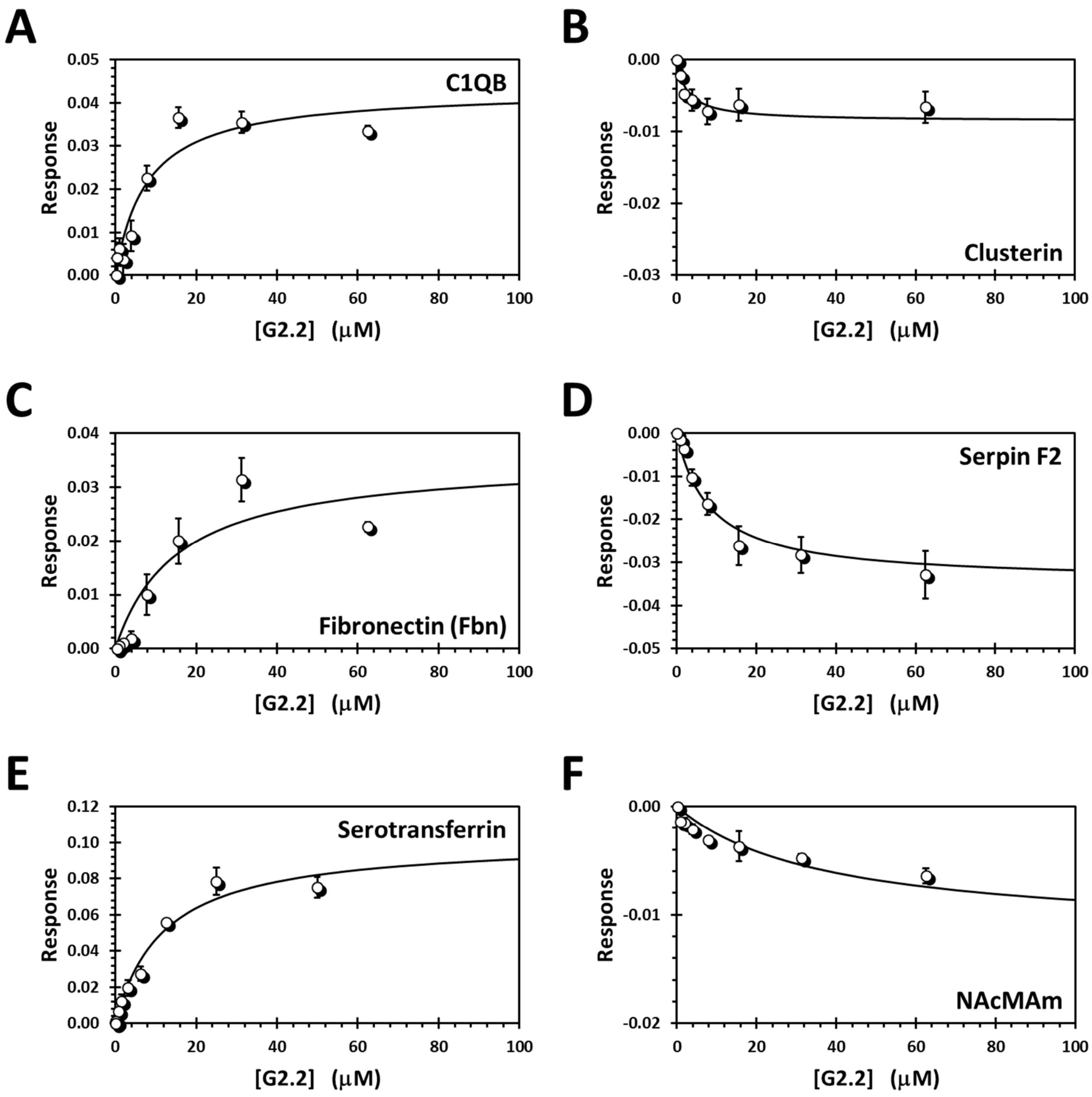


**Figure S6.** Binding affinity of G2.2 for the six representative protein targets identified through PAL-Pomics. His-tagged proteins, labeled with RED-tris-NTA dye, were used in microscale thermophoresis (MST)-based titrations to measure $K_D$ of interaction. Solid line shows non-linear regression using the standard hyperbolic binding equation to calculate the $K_D$.
